# Structural basis for loss of covalent flavinylation in the H158Y mutant of pyranose oxidase

**DOI:** 10.64898/2026.02.17.706288

**Authors:** Yuki Yashima, Clemens Karl Peterbauer, Taku Uchiyama, Kota Takeda, Kiyohiko Igarashi

**Author notes:** **Correspondence:** K. Igarashi: Forest Chemistry Laboratory, Department of Biomaterial Sciences, Graduate School of Agricultural and Life Sciences, The University of Tokyo, Yayoi, Bunkyo-ku, Tokyo 113–8657, Japan, (should not be published.).

## Abstract

Pyranose oxidase from *Phanerochaete chrysosporium* (*Pc*POx) is a flavoprotein that forms a covalent 8α-*N*^3^-histidyl-FAD linkage at His158. Here, we determined the crystal structure of the *Pc*POx H158Y variant at 1.69 Å resolution and investigated its FAD occupancy and catalytic activity. Tyr158 adopts a rotamer conformation incompatible with covalent flavinylation, placing its phenolic Oη atom 8.7 Å away from the FAD C8α atom. This conformation is stabilized through a Tyr158–Lys79 hydrogen bond. The additional mutation K79A abolishes this hydrogen bond, potentially freeing the Tyr158 residue for other conformations, but does not restore covalent flavin attachment. The H158Y variant retains substantial FAD occupancy and tetrameric assembly but shows markedly reduced oxidase and dehydrogenase turnover rates (less than 13% of WT). These results provide a structural explanation for the failure of tyrosine substitution to support covalent flavinylation in *Pc*POx and offer insights into requirements for engineering alternative covalent flavin linkages.

## INTRODUCTION

Flavin cofactors are among the most common prosthetic groups found in oxidoreductases. Some flavin-dependent enzymes form covalent bonds between their amino acid residues and the flavin cofactors, either FAD (flavin adenine dinucleotide) or FMN (flavin mononucleotide). To date, several types of monocovalent linkages between amino acid residues and the isoalloxazine ring of flavins have been identified, including 8α-*N*^1^-histidyl-FAD [1], 8α-*N*^3^-histidyl-FAD [2], 8α-*S*-cysteinyl-FAD [3], 8α-*O*-tyrosyl-FAD [4], 8α-*O*-aspartyl-FAD [5], 6-*S*-cysteinyl-FAD [6], and 6-*S*-cysteinyl-FMN [7]. In addition, a bicovalent flavin linkage involving 8α-*N*^1^-histidyl-FAD and 6-*S*-cysteinyl-FAD has been reported [8]. FMN derivatives linked to threonine or serine residues via phosphoester bonds are known as phosphoester flavinylations [9,10].

These covalent flavin attachments play crucial roles in regulating enzymatic activity, stability, and redox potential [11]. Accordingly, engineering artificial covalent flavin attachment in proteins with noncovalently bound flavin has attracted increasing interest. Previous studies have achieved covalent attachment of chemically modified flavins to enzymes [12–15], and there have also been successful examples of introducing covalent bonds between enzyme variants and FAD or FMN [16,17]. Some of these engineered flavoproteins exhibited higher redox potentials [16] or enhanced catalytic activity and thermostability [17]. However, our understanding of how to generate artificial covalent flavins remains limited, particularly with respect to switching covalent linkage patterns in naturally covalent flavoproteins.

The monocovalent flavoprotein pyranose 2-oxidase (POx; EC 1.1.3.10) contains an 8α-*N*^3^-histidyl-FAD linkage [2] (Fig. 1A&B). POx catalyzes the oxidation of the C2-hydroxyl group of aldopyranoses, accompanied by reduction of molecular oxygen to hydrogen peroxide [18]. In addition, POx can reduce quinones, phenoxy radicals, and metal ions [18–20]. Owing to these properties, its physiological role has been proposed to involve hydrogen peroxide generation and reduction of phenoxy radicals or metal-ion during lignin degradation in fungi [20–22] and bacteria [19]. POx exhibits regioselectivity toward sugar substrates; it generally oxidizes the C2-hydroxyl group, but can also oxidize the C3-hydroxyl group of 2-deoxy-D-glucose [23].

**Fig. 1.**
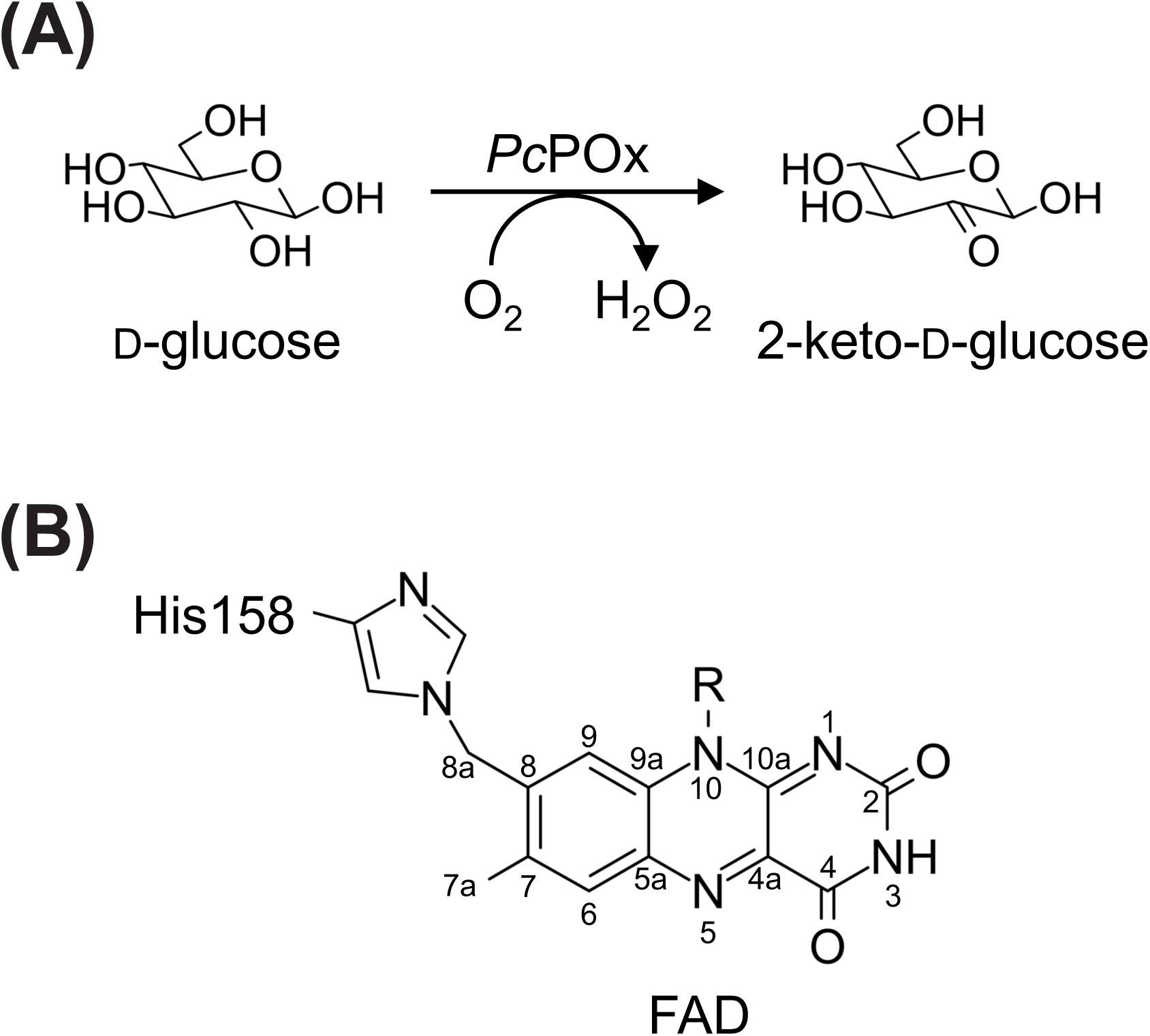
Reaction of *Pc*POx (A) and covalent FAD in *Pc*POx (B). The chemical structures were drawn using Marvin Sketch (Chemaxon Ltd., Budapest, Hungary).

The covalent linkage of FAD in POx has long been of interest. Substitution of the histidine residue responsible for covalent FAD attachment with alanine results in noncovalent FAD binding in POx from *Trametes ochracea* (heterotypic synonym: *Trametes multicolor*) (*To*POx) [24] and *Phanerochaete chrysosporium* (*Pc*POx) [25]. In particular, the *To*POx H167A mutant exhibits decreased catalytic turnover and redox potential compared with the wild-type enzyme [24]. Recently, we investigated the potential to introduce alternative covalent flavinylation patterns in *Pc*POx by comparing 19 variants at His158, which is responsible for the covalent FAD attachment [26]. None of the His158 variants formed covalent flavin bonds, although 17 of 19 retained FAD noncovalently. Two variants, H158D and H158P, produced apoenzymes lacking FAD. Nevertheless, it remained unclear why tyrosine and cysteine, which can form covalent flavin linkages in other enzymes, failed to do so in *Pc*POx. Structural information explaining why these substitutions fail to support covalent flavinylation has been lacking.

In this study, we determined the crystal structure of the *Pc*POx H158Y variant at 1.69 Å resolution and characterized its FAD occupancy and catalytic properties to clarify why tyrosine substitution does not support covalent flavinylation in *Pc*POx. The relationship between the H158Y mutation and the enzymatic properties of *Pc*POx is discussed.

## MATERIALS AND METHODS

### Chemicals and strains

All chemicals were purchased from FUJIFILM Wako Pure Chemical Corp. (Osaka, Japan) except for LB Broth and PEG3350 from Sigma-Aldrich, and *N*-ethyl-*N*-(2-hydroxy-3-sulfopropyl)-3-methylaniline, sodium salt (TOOS) from DOJINDO LABORATORIES Corp. (Kumamoto, Japan). For recombinant enzyme expression, *ECOS*^TM^ SONIC Competent *E. coli* strain BL21 (DE3) containing a plasmid with pET-27b (+) vector and *Pc*POx WT or H158Y gene was prepared in the previous work [26].

### Cultivation

For production of *Pc*POx enzymes applied for biochemical analysis, a selected single colony of *E. coli* containing the respective expression plasmid was inoculated into 5 mL liquid LB medium containing 50 μg·mL^−1^ kanamycin and shaken at 37 °C and 300 rpm overnight. 4 mL of the preculture was added to 200 mL liquid TB medium without antibiotics and shaken (150 rpm) at 37 °C for 2 hours. *Pc*POx gene expression was induced with 0.5% (w/v) lactose overnight at 26.5 °C and 150 rpm. Cells were collected by centrifugation (3,000 rcf, 8 °C, and 5 min). The cell pellet was washed with approximately 100 mL of pure water and cells were collected by centrifugation again. The cell pellet was re-suspended in 20 mL of binding buffer; 20 mM sodium phosphate (pH 7.4) containing 300 mM NaCl and 50 mM imidazole. The re-suspended cells were disrupted in an ultrasonic homogenizer (VP-300N, TAITEC Corporation, Saitama, Japan). The homogenate was centrifuged (15,000 rcf, 8 °C, 30 min), and the supernatant was collected. The cell extract solution was filtered through a syringe filter (Millex-GV Syringe Filter Unit, 0.22 μm pore size, PVDF, gamma sterilized; Merck KGaA). For the enzyme production of *Pc*POx H158Y for crystallization, the cultivation was scaled up to 2 L LB medium. Recombinant expression was induced by 0.4 mM IPTG and continued overnight at 26.5 °C.

### Purification

Crude enzyme solution was applied to a 1 mL HisTrap^TM^ HP column (Cytiva, Tokyo, Japan) equilibrated with the binding buffer described above. The flow rate was set to 1 mL·min^−1^ for this purification step. After washing the column with 15-20 column volumes (CV) of binding buffer, bound histidine-tagged proteins were eluted by 8 CV gradient elution with 0-100 % elution buffer; 20 mM sodium phosphate (pH 7.4) containing 300 mM NaCl and 500 mM imidazole. Yellow fractions containing oxidase activity toward glucose were collected. Protein solutions were buffer-exchanged to 100 mM sodium chloride in 50 mM sodium phosphate buffer (pH 7.0) using VivaspinTM 20-10K (Cytiva) for storage. For enzyme production of *Pc*POx H158Y for crystallization, a 5 mL HisTrap^TM^ HP column (Cytiva) was applied for the first purification step. To further improve purity, anion exchange chromatography as a second purification step was conducted. Protein solution was buffer-exchanged to 20 mM sodium phosphate buffer (pH 6.5) using ultrafiltration discs (5 kDa NMW; Merck KGaA). Protein solution was applied to TSKgel DEAE-5PW (21.5 mm × 15 cm) equilibrated with 20 mM sodium phosphate buffer (pH 6.5). The flow rate was set to 4 mL・min^−1^ for this purification step. After washing with 8 CV equilibration buffer, elution was performed by an 8 CV gradient (0-200 mM sodium chloride in 20 mM sodium phosphate buffer, pH 6.5). Yellow fractions containing oxidase activity toward glucose were collected.

### Protein concentrations

Protein concentrations were determined by measuring absorbance at 280 nm using a NanoDrop^TM^ 2000 spectrophotometer (Thermo Fisher Scientific, Waltham, MA, USA). The extinction coefficients at 280 nm were calculated using the online tool ProtParam [27] (https://web.expasy.org/protparam/) accessed on 22 September 2025 and 9 January 2026, and were 0.965 mL·mg^−1^cm^−1^ for *Pc*POx WT, 0.986 mL·mg^−1^cm^−1^ for *Pc*POx H158Y, and 0.986 mL·mg^−1^cm^−1^ for *Pc*POx H158Y/K79A. The molecular weights of all *Pc*POx enzymes were calculated to be 70 kDa using ProtParam.

### Protein crystallography

Purified *Pc*POx H158Y variant was buffer-exchanged and concentrated to 32 mg·mL^−1^ in 200 mM HEPES buffer pH 7.2 using Vivaspin^TM^ 500-10K (Cytiva). Screening of protein crystallization conditions was conducted using a screening kit, NeXtal JCSG + Suite (Qiagen, Hilden, Germany). Finally, 3 µL protein sample solution and 3 µL reservoir solution (B2 of JCSG + Suite; 0.2 M sodium thiocyanate, 20% (w/v) PEG3,350) were mixed using the sitting-drop vapor-diffusion method and incubated at 293 K for a month. For data collection, the *Pc*POx H158Y crystal was incubated in cryoprotectant solution containing 0.1 M sodium thiocyanate, and 20% (w/v) PEG3,350, 25% (w/v) glycerol for a few minutes and then flash-frozen in liquid nitrogen.

X-ray diffraction dataset of *Pc*POx H158Y crystal was collected using synchrotron radiation at the beamline BL-5A of the Photon Factory (Tsukuba, Japan). Diffraction images were converted to XDS format using XDS [28] integrated into PReMo [29]. Scaling and merging were carried out with AIMLESS [30] in CCP4 (v.8.0.19). Molecular replacement was performed with an AlphaFold 3 [31] -predicted model using Molrep in CCP4. The predicted model contained FAD and full length of *Pc*POx H158Y with a histidine tag as a monomer. Refinement was performed initially with Refmac (v. 5.8.0425) [32] in CCP4, and finalized with phenix.refine in PHENIX (v.1.21.2) and COOT (v. 9.8.93) [33]. The structural graphics were drawn with PyMOL (v.3.1.6.1).

### Electrophoresis

SDS-PAGE analysis using 12% polyacrylamide gel and the detection of covalent FAD were conducted as described in the previous work [26]. Native-PAGE analysis was conducted with 0.34 M Glycine, 0.265 M Tris-HCl buffer pH 8.9 for the cathode, and 0.5 M Tris-HCl buffer pH 7.8 for the anode. Protein samples were diluted with ×2 sample buffer: 125 mM Tris-HCl pH 6.8, 20% Glycerol, 0.01% BPB(bromophenol blue). Samples were loaded into 7.5 % polyacrylamide gel and run at 100 V for 4 hours. The protein bands were visualized using SimplyBlue^TM^ SafeStain (Invitrogen, Carlsbad, CA, USA).

### UV-visible spectra

Ultraviolet-Visible (UV) -visible spectra of 35.7 μM (2.5 mg·mL^−1^) *Pc*POxs in 100 mM sodium chloride in 50 mM sodium phosphate buffer (pH 7.0) were recorded at room temperature (26.5 ℃) using a Jasco V660 spectrophotometer (Jasco Ltd., Tokyo, Japan). Spectra were recorded at a scan rate of 100 nm·min^−1^, with a data interval of 0.1 nm, over the range 250–700 nm. Trichloroacetic acid (TCA) precipitation was performed as described by Macheroux [34] with the following modifications: denatured protein was precipitated by centrifugation at 20,000 rcf for 10 mins, and we did not add solid sodium carbonate to adjust pH as in the previous work [35]. FAD occupancy of *Pc*POx H158Y was calculated using released FAD (ε450 = 11.3 mM^−1^cm^−1^) spectra after TCA precipitation. FAD occupancy of *Pc*POx WT was estimated from the *R*z ratio (Absorbance at a peak derived from FAD around 450 nm / Absorbance at 280 nm) and the FAD occupancy of *Pc*POx H158Y, determined by the equation below (1). The spectra were drawn using Igor Pro 9 ver. 9.5 (WaveMetrics, Inc., Portland, OR, USA).

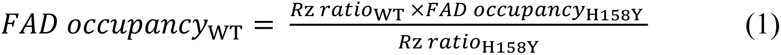

### Activity assay and steady-state kinetic analysis

All activity assays were conducted using 96-well plates and a Multiskan SkyHigh Microplate Spectrophotometer (ThermoFisher) without oxygen control. Reactions were monitored for 5 min (in oxidase assay) or 3 min (in dehydrogenase assay) at 26.5 ℃ in 50 mM sodium phosphate buffer pH 7.0. Steady-state kinetic parameters were calculated by GraphPad Prism version 10.5.0 (GraphPad Software Inc., California, US) using the Michaelis-Menten equation.

Oxidase activity was measured using a horseradish peroxidase assay reported previously [26]. Enzyme concentration in a final reaction mixture was 35 nM for *Pc*POx WT and 140 nM *Pc*POx variant H158Y (700 nM H158Y only for galactose and 2-deoxy-D-glucose). For kinetic analysis, 0.1-20 mM glucose, 2-200 mM xylose, 1-100 mM galactose, and 10-500 mM 2-deoxy-D-glucose were tested for *Pc*POx WT. Due to the difference in *K*_m_ values, 1-200 mM glucose, 5-400 mM xylose, 2-200 mM galactose, and 30-500 mM 2-deoxy-D-glucose were tested for *Pc*POx variant H158Y. At least 7 substrate concentrations under ambient oxygen conditions were tested to obtain kinetic parameters.

For dehydrogenase activity, 2,6-dichloroindophenol (DCIP) and 1,4-benzoquinone (BQ), were used in the present of ambient oxygen. The reaction mixture contained 50 mM glucose, 50 mM sodium phosphate buffer pH 7.0, enzyme (100 nM *Pc*POx WT or 500 nM *Pc*POx H158Y), 500 µM electron acceptor: DCIP (ε_520_ = 6.6 mM^−1^cm^−1^), and BQ (ε_290_ = 2.24 mM^−1^cm^−1^).

### Prediction of p*K*_a_ values of amino acid residues in crystal structures

The p*K*_a_ values of amino acid residues in crystal structures were predicted using an online tool, PropKa (https://www.ddl.unimi.it/vegaol/propka.htm), accessed on 1 January 2026 [36].

### Site-directed mutagenesis

Overlap extension PCR [37] was employed for site-directed mutagenesis of H158Y/K79A in *Pc*POx, using the expression plasmid as template. The nucleotide sequences of the synthetic primers are given in Table S1. DNA polymerase KOD One^®^ PCR Master Mix (Toyobo Co., Ltd., Osaka, Japan) was used. The amplified product was transformed into *ECOS*^TM^ SONIC Competent *E. coli* strain BL21 (DE3) by heat-shock. The single colony containing the correct plasmid was used for the protein expression as described above.

## RESULTS

### Sequence differences in *Pc*POx are unlikely to affect the active site or FAD environment

The amino acid sequence of *Pc*POx we used in this research contained two amino acid substitutions (R9S and K509N) compared to the deposited sequence in the previous work (PDB ID 4MIF) although both *Pc*POxs were cloned from the same strain K-3 [25]. The mutation R9S is in the flexible N-terminus loop and the K509 is on the surface of the enzyme (Fig. S1A,B). They are far from the FAD cofactor and the residue His158. There were 13 mutations in the nucleic acid sequences in our *Pc*POx, and 12 of them were located in the third base in codons (Table S1). All experiments described below were performed *Pc*POx containing Ser9 and Asn509 as the *Pc*POx WT.

### H158Y retains FAD occupancy and tetrameric assembly

To assess the biological assembly of the variant H158Y, electrophoresis was carried out. In SDS-PAGE analysis, *Pc*POx H158Y showed a band at the same location as the WT, and did not exhibit any fluorescence derived from the covalently bound flavin (Fig. 2A,B). Native-PAGE analysis indicated the mutation of H158Y did not affect the quaternary structure (Fig. 2 C). These results demonstrated the variant H158Y with noncovalently bound FAD was expressed successfully as a full-length polypeptide and intact tetramer structure like the WT [38, 39].

**Fig. 2.**
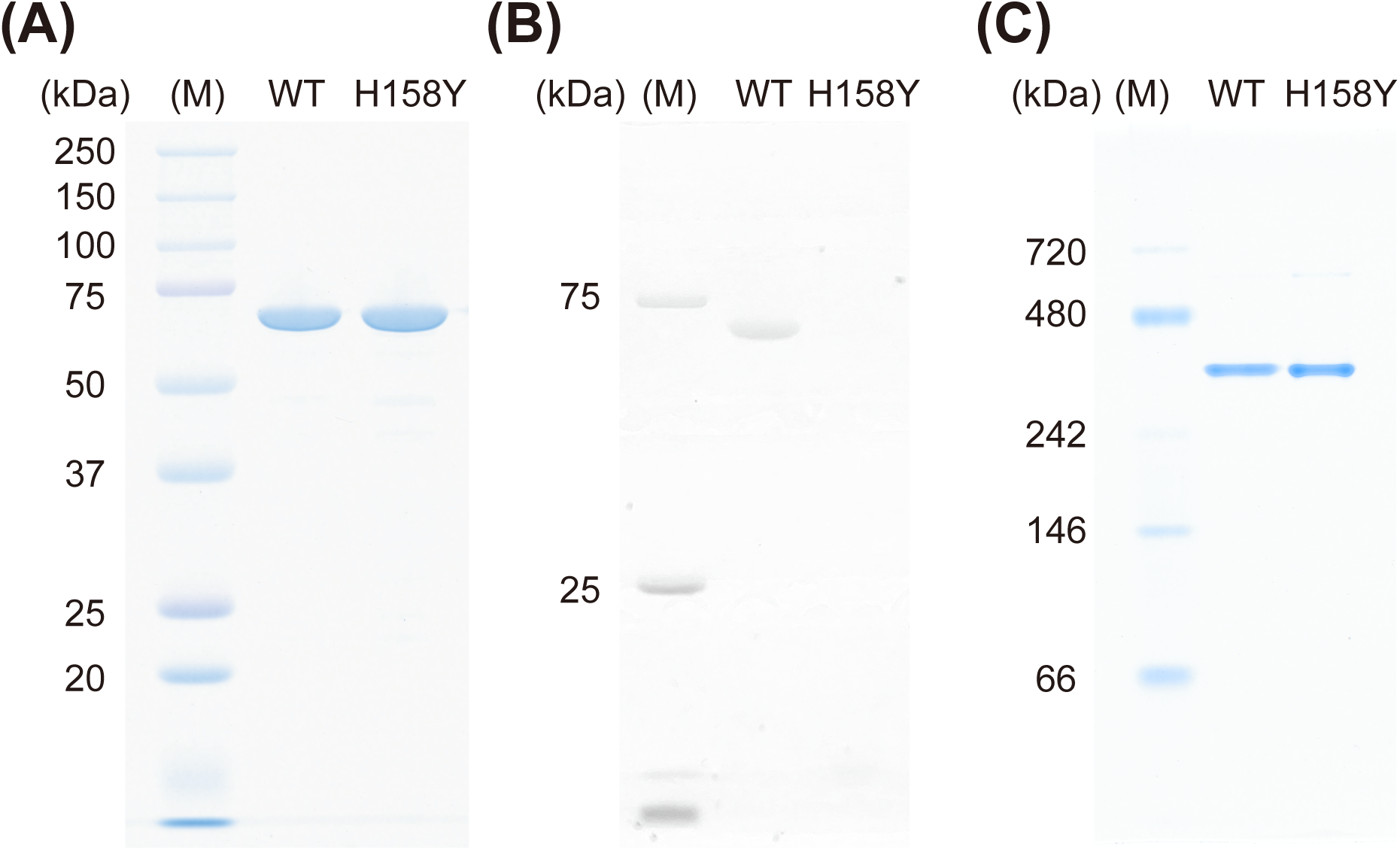
Electrophoresis of *Pc*POx WT and H158Y. (A) SDS-PAGE analysis, (B) Detection of covalently bound FAD under UV illumination (365 nm), and (C) Native-PAGE analysis.

To estimate FAD occupancy of *Pc*POx WT and the variant H158Y, UV-visible spectra analysis was conducted (Fig. 3 AB). *Pc*POx WT exhibited the two peaks at 361.3 nm and 459.3 nm derived from the FAD cofactor, while the variant H158Y showed the two peaks at 383.2 nm and 451.4 nm. The blue shift seen in the spectra of WT (361.3 nm) is attributable to the covalent FAD. Although the *R*z ratio (Absorbance at a peak derived from FAD around 450 nm / Absorbance at 280 nm) was typical for the analysis of flavin-dependent protein, the reciprocal of it (1/ *R*z) was commonly used for covalent flavoprotein [26, 40]. *Pc*POx WT and the variant H158Y exhibited almost the same 1/*R*z values (12.8 for WT and 12.4 for H158Y). Through TCA precipitation, the FAD occupancy of the H158Y variant was determined to be 46.9%. In comparison with the 1/*R*z values for WT and H158Y, the FAD occupancy of the WT was calculated as 45.4%.

**Fig. 3.**
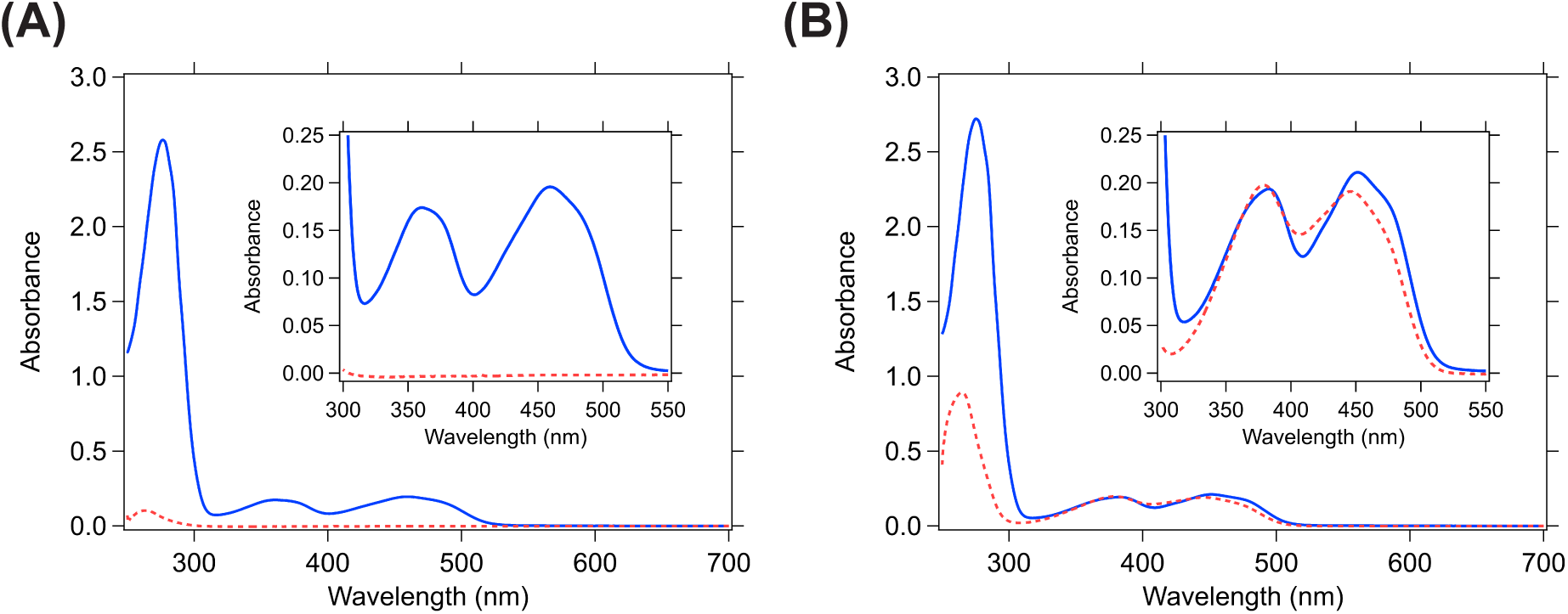
UV-visible spectra of 35.7 µM *Pc*POx WT (A) and H158Y (B) before (blue solid line) and after (red dotted line) TCA precipitation. Expanded view from 300 to 550 nm are shown in the inset.

### The H158Y mutation does not alter the overall fold of *Pc*POx

Using the 2-step purified *Pc*POx H158Y sample, the yellow, rectangular plate-like crystals were obtained. X-ray crystallographic data collection and refinement statistics are summarized in Table 1. The structure of *Pc*POx H158Y was solved at the maximum resolution of 1.69 Å. The *R*_work_ value (0.2131) and the *R*_free_ value (0.2404) were relatively high considering the resolution due to the missing residues, which was almost corresponding to the missing regions of recombinant *Pc*POx WT (PDB ID: 4MIG) [25]. The disordered regions were the N-terminus (residues1-12), a flexible loop (residues 57-69), Leu131, a flexible loop (residues 309-318 and residues 349-365), the substrate recognition loop (residues 459-465) and the C-terminus (downstream from residue 617). The RMSD value compared with chain A of the recombinant *Pc*POx WT (PDB ID: 4MIG) was 0.191 Å and no significant overall structural change was observed. Although the space group *P*4_2_2_1_2 was different from the recombinant *Pc*POx WT and H158A (*P*12_1_1), *P*4_2_2_1_2 is common for some variants of *To*POx (PDB ID: 3PL8, 4MOF, 4MOG, 4MOH, 4MOI, 4MOO, and 4MOS) [41][42]. As well as those structures, the asymmetric unit contained one polypeptide.

**Table 1.**
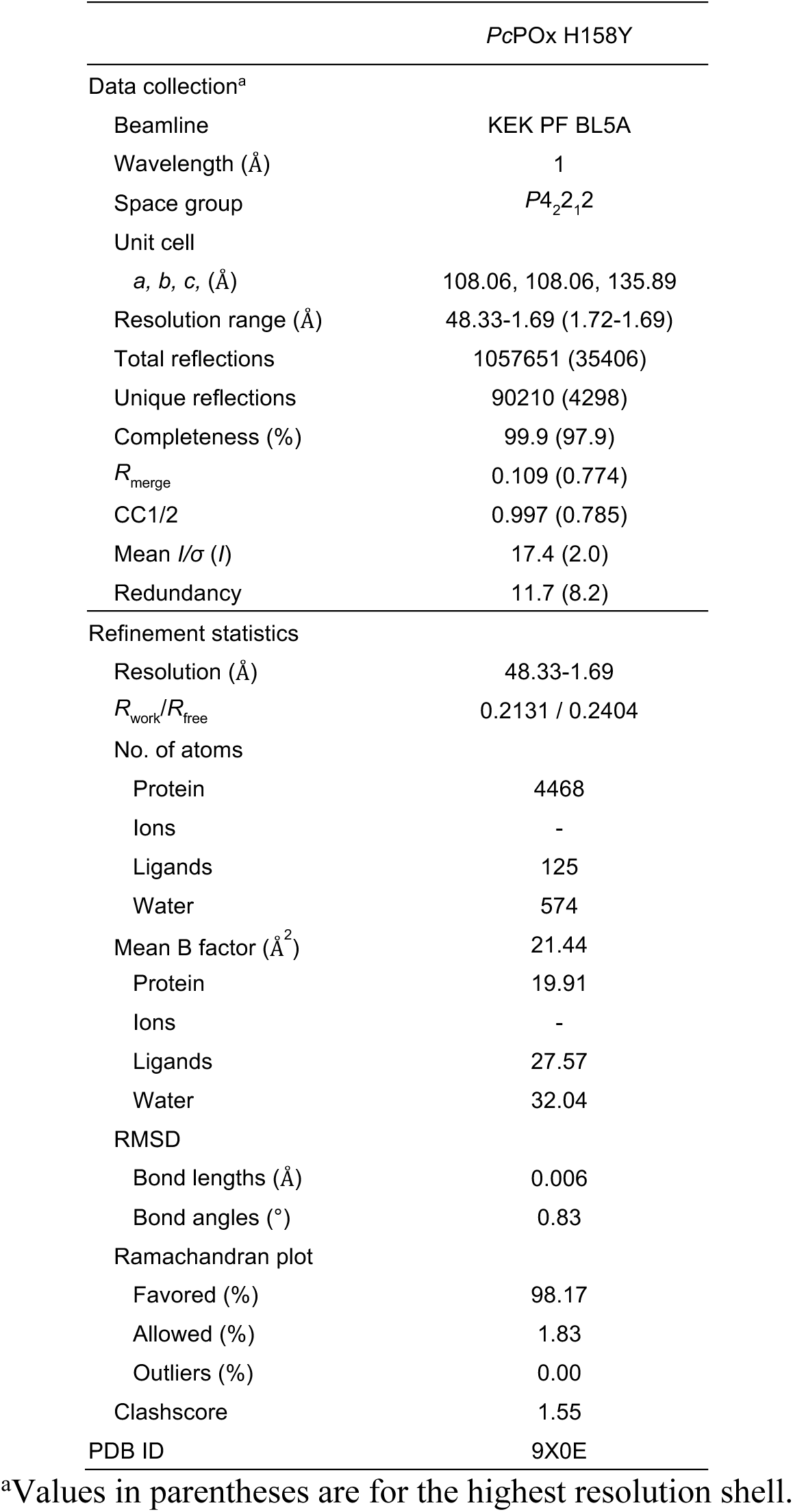
Statistics of crystallographic data collection and refinement.

### The active-site geometry is conserved except for residue 158

The catalytic centre in *Pc*POx H158Y mutant is shown in Fig. 4 A. The side chain of Tyr158 points in an inverse direction away from C8α of FAD. Still most of the amino acid residues in the catalytic centre except for Tyr158 did not show any conformational change compared to *Pc*POx WT (PDB: 4MIG) and the H158A mutant (PDB: 4MIH) [25] including the catalytic pair residues His553 and Asn596 (Fig. S2). The structure of *Pc*POx H158Y contains a glycerol molecule in the active site (Fig. 4 B). Although we tried soaking experiments with 3-deoxy-3-fluoro-D-glucose and mannose as slow-rate substrates, we could not obtain diffraction data of sufficient quality. The secondary hydroxyl group of glycerol forms hydrogen bonds with Nε2 of His553 (2.9 Å), Nδ2 of Asn596 (2.8 Å), and N5 of FAD (3.3 Å). This orientation of the glycerol is consistent with that of sugar substrates in *Pc*POx WT (Fig. 4 C). The substrate recognition loop (^458^DAFSYG^463^) did not exhibit sufficient electron density for modeling except for Asp458. The carboxylate oxygens (Oδ1 and Oδ2) of Asp458 form hydrogen bonds to the O γ1 atom of Thr160 (2.7 Å and 3.4 Å) (Fig. S3).

**Fig. 4.**
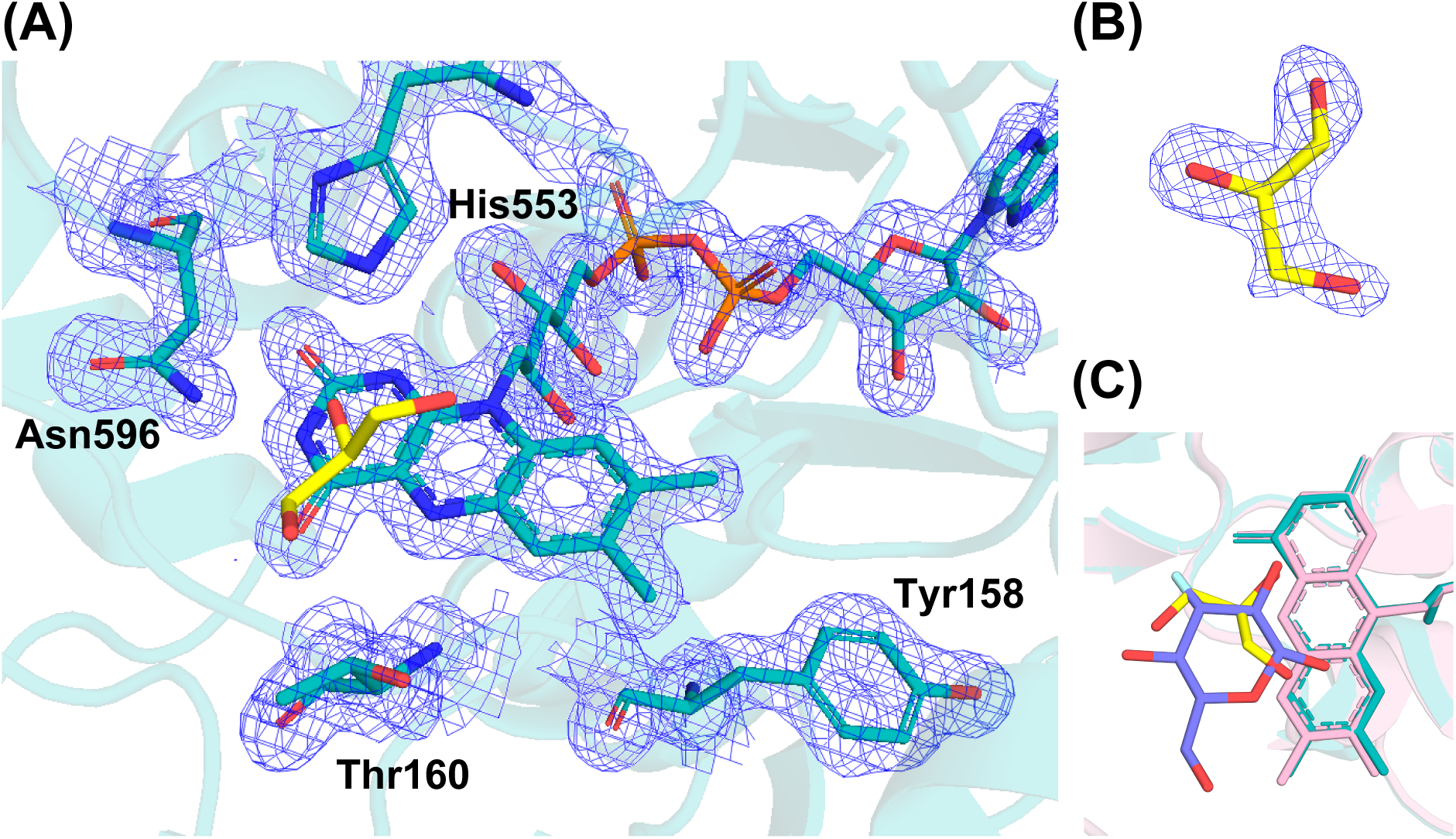
Crystal structure of *Pc*POx H158Y mutant. (A) Active site of *Pc*POx H158Y. (B) Electron density map of the glycerol molecule in the active site of the *Pc*POx H158Y mutant. (C) Comparison of the orientation of the substrate analog 3-deoxy-3-fluoro-D-glucose (purple) in *Pc*POx WT (PDB ID: 4MIG) and the glycerol molecule (yellow) in the H158Y mutant. The structures were aligned by pair fitting of FAD cofactors. The protein backbone and FAD cofactor are shown in cyan (H158Y) and magenta (WT). The 2*F*_o_−*F*_c_ electron density map is contoured at 1.5 σ.

### Tyr158 adopts a rotamer incompatible with C8α covalent flavin linkage and is not rescued by K79A

Loss of the covalent flavin linkage resulted in a flatter isoalloxazine ring, as observed in the H158A mutant, causing the C8α atom moving for 0.9 Å to become a flat ring. This flattening of the flavin isoalloxazine ring also happened in the *Pc*POx mutant H158A (PDB: 4MIH) [25].

The side chain of Tyr158 adopted an altered rotameric conformation relative to the original histidine residue (Fig. 5A). As a result of the conformational change, the distance between Oη of Tyr158 and C8α of FAD was 8.7 Å, too far to form a covalent interaction. The conformational change of Tyr158 causes a hydrogen bond (3.0 Å) with Oη of Tyr158 and Nζ of Lys79 to form. This interaction made the pKa value of the side chain of Tyr158 decrease to 8.96 in the structure-based prediction (Table S3). The double mutant K79A/H158Y was prepared to eliminate fixing the unfavorable Tyr158 conformation by Lys79 (Fig. 5B-D). As in the 158Y mutant, the K79A/H158Y mutant did not form a covalent interaction with FAD. In UV-vis spectra, the K79A/H158Y mutant showed peaks at 384.5 nm and 451.0 nm, highly similar to the H158Y mutant. The 1/*R*z value was 10.0, and FAD occupancy was 64.8 % for the H158Y/K79A mutant.

**Fig. 5.**
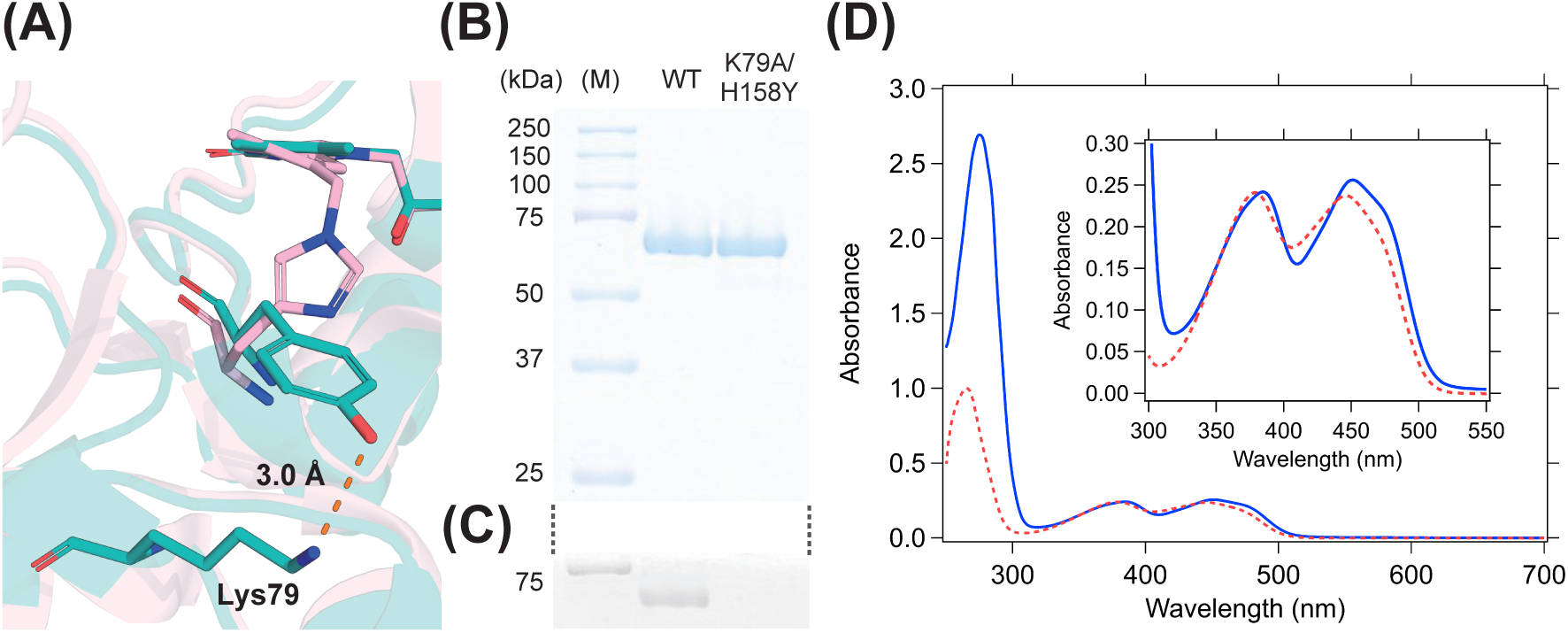
Interaction between Tyr158 and Lys79 in *Pc*POx H158Y mutant. (A) Conformation of Tyr158 in *Pc*POx H158Y mutant, compared with His158 in *Pc*POx WT. Coloring for *Pc*POx WT (PDB ID: 4MIG) and the variant H158Y is the same as Fig. 4. (B) SDS-PAGE analysis of *Pc*POx variant K79A/H158Y. (C) Detection of the covalent flavin cofactor using the same gel as (B). (D) UV-visible spectra of 35.7 µM *Pc*POx variant K79A/H158Y before (blue solid line) and after (red dotted line) TCA precipitation. Expanded view from 300 to 550 nm are shown in the inset.

**Fig. 6.**
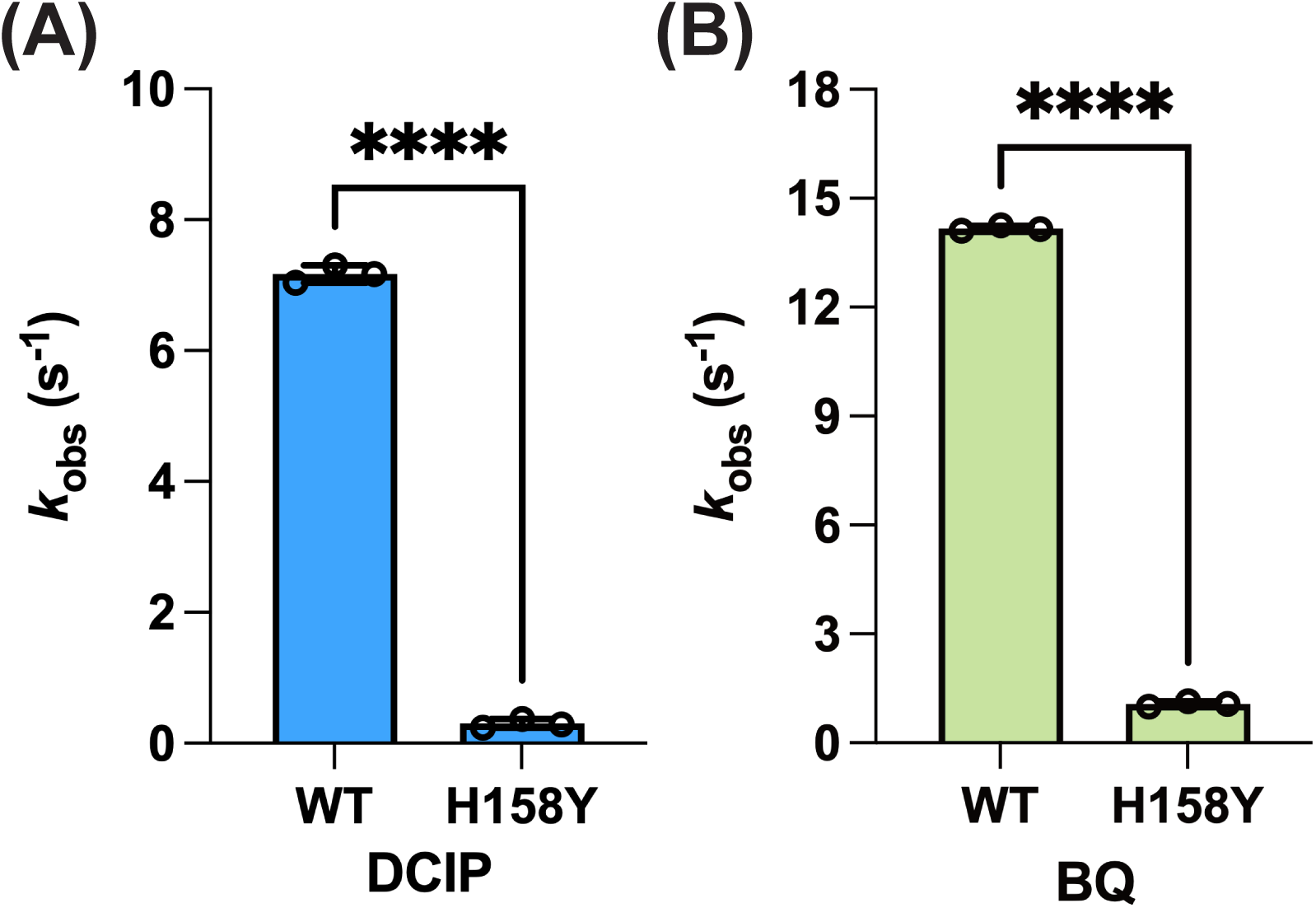
Dehydrogenase activity of *Pc*POx WT and H158Y in 50 mM sodium phosphate buffer (pH 7.0). Considering the *K*_m_ values for glucose in the enzymes, 50 mM glucose was used as the electron donor, and 500 µM DCIP (A) or BQ (B) was used as the electron acceptor. Bars represent mean ± standard deviation (SD) of three technical replicates (n = 3). Circles represent individual measurements. Statistical significance was evaluated using Welch’s *t*-test (****, *p*<0.0001).

### Loss of covalent flavinylation severely reduces oxidase and dehydrogenase turnover

To understand the effect of the mutation of H158Y on enzymatic activity, steady-state kinetic constants for various sugar substrates were determined using the horseradish peroxidase assay (Table 2). The curve-fittings of Michaelis-Menten equation were given in Fig. S4. Since POx catalyzes two typical oxidation modes, C2-oxidation and C3-oxidation, we tried glucose, xylose, and galactose for C2-oxidation and 2-deoxy-D-glucose for C3-oxidation. The enzymatic activitiy of *Pc*POx H158Y decreased significantly for all tested sugar substrates including C2 and C3-oxidation modes (less than 13% of *k*_cat_ values, and 7% of *k*_cat_/*K*_m_ values of the WT) although the *K*_m_ values differed by sugar substrates. We evaluated not only the oxidase activity but also the dehydrogenase activity under ambient oxygen. The *k*_obs_ values (turnovers) of dehydrogenase activity using 500 µM DCIP or BQ as an artificial electron acceptor (Fig. 4) were compared since steady-state kinetic analysis toward electron acceptors was difficult to measure due to oxygen inhibition. The *k*_obs_ values in the DCIP assay was 7.2 s^−1^ (WT) and 0.3 s^−1^ (H158Y). The *k*_obs_ values in the BQ assay was 14 s^−1^ (WT) and 1.1 s^−1^(H158Y). In both conditions, the mutation H158Y decreased the dehydrogenase activity to less than 8% compared to WT.

**Table 2.** Steady-state kinetic parameters of oxidase activity toward various sugar substrates in *Pc*POx WT and the variant H158Y (pH 7.0). The symbol ± indicates the standard error of three technical replicates.

|  |  | WT |  |  | H158Y |  |  |
| --- | --- | --- | --- | --- | --- | --- | --- |
| Substrate (oxidation mode) | | $K_M$ (mM) | $k_{cat}$ ($s^{-1}$) | $k_{cat}/K_M$ ($s^{-1}mM^{-1}$) | $K_M$ (mM) | $k_{cat}$ ($s^{-1}$) | $k_{cat}/K_M$ ($s^{-1}mM^{-1}$) |
| Glucose | (C2) | $0.88 \pm 0.04$ | $9.3 \pm 0.1$ | 11 | $3.7 \pm 0.2$ | $1.1 \pm 0.0$ | 0.31 |
| Xylose | (C2) | $20 \pm 1$ | $5.0 \pm 0.1$ | 0.25 | $25 \pm 1$ | $0.32 \pm 0.00$ | 0.012 |
| Galactose | (C2) | $9.5 \pm 0.5$ | $0.59 \pm 0.01$ | 0.062 | $18 \pm 3$ | $0.076 \pm 0.002$ | 0.0042 |
| 2-Deoxy-D-glucose | (C3) | $47 \pm 4$ | $1.4 \pm 0.0$ | 0.033 | $58 \pm 11$ | $0.10 \pm 0.05$ | 0.0018 |

## DISCUSSION

The covalent interaction between flavin cofactors and amino acid residues in flavoproteins has been well studied, using crystal structures of alanine mutants at the covalent flavinylation sites [24, 40, 43]. However, three-dimension structures of mutants carrying alternative nucleophilic residues (e.g., aspartic acid, cysteine, and tyrosine) have rarely been reported, leaving the structural basis for the failure of covalent flavinylation in the mutants unclear. The aim of this research is to elucidate the structural reason of incapability of forming a covalent interaction to the FAD cofactor in *Pc*POx H158Y mutant and to assess the impact of the loss of covalent flavinylation on FAD properties and catalytic activity. The crystal structure of *Pc*POx H158Y revealed that Tyr158 adopts a rotamer conformation that is incompatible with an 8α covalent bond to FAD. The phenolic oxygen of Tyr158 is positioned 8.7 Å away from the FAD C8α atom, which far exceeds the distance compatible with covalent bond formation. In addition, Tyr158 forms a hydrogen bond with Lys79, which may stabilize this unproductive Tyr158 conformation. To test this hypothesis, we generated the H158Y/K79A double mutant to disrupt the Tyr158–Lys79 hydrogen bond. However, covalent flavinylation was not restored in this variant, indicating that removal of the Tyr158–Lys79 interaction is not sufficient to enable formation of an 8α-*O*-tyrosyl-FAD linkage in *Pc*POx. This suggests that additional structural constraints around the flavinylation site prevent productive positioning of Tyr158 for covalent bond formation.

Our *Pc*POx contains two amino acid substitutions (R9S and K509N) relative to the previously deposited *Pc*POx sequence (PDB ID:4MIG), although both sequences were cloned from the same strain K-3 [25,26,44]. On the nucleic acid level, 12 of 13 substitutions were located in the third base of codons, and 11 of these were silent sequence variation. Considering *Phanerochaete chrysosporium* K-3 strain is a heterokaryon [45,46], this nuleic acid substitution patterns suggested that the *Pc*POx we cloned and used in this study was an allozyme of the *Pc*POx in the original study. The two amino acid substitutions are located far from the catalytic centre and the FAD cofactor including the histidine158. Therefore, they are unlikely to affect the active-site geometry or the conclusions drawn from the structural comparisons among *Pc*POx WT (PDB ID: 4MIG), H158A (PDB ID: 4MIH), and H158Y (this study).

*Pc*POx enzymes exhibited 45-65 % of the FAD occupancy estimated by TCA precipitation and the 1/*R*z value. Although the cause of the presence of apoenzyme remains unclear, these values are consistent with those reported for other flavoproteins. For example, in the flavoprotein vanillyl-alcohol oxidase, where the flavin is covalently linked to the protein, the A280/A439 ratio of the WT is 13.4 [47], whereas the corresponding ratio of the variants H422A, H422C, and H422T, which show noncovalent flavin attachment, is 11.5 [40]. In case of pyranose oxidase from *Oscillatoria princeps* (*Op*POx), the A280/A450 ratio is 11.7 [48]. These ratio is comparable to the 1/*R*z value obtained for *Pc*POx enzymes. Determination of FAD occupancy by TCA precipitation is widely used and has yielded occupancies ranging from 9 % to 70 % for various recombinant cellobiose dehydrogenases [49–51]. Taken together, our results suggest that the H158Y mutation does not substantially affect FAD occupancy compared with the WT of recombinant *Pc*POx.

POx exhibits mainly three conformational states; closed mode, semi-open mode, and open mode, which are coupled to conformational changes of the substrate recognition loop [24,25,41,52]. The closed conformation binding acetate in *To*POx is considered the conformation for the oxidative half-reaction, which is a state suitable for stabilization of a C4a-hydroperoxyflavin [52, 53]. The semi-open mode is compatible with C2-oxidation [41], whereas the open mode is associated with C3-oxidation mode [24]. Although we could not model most residues of the substrate recognition loop ^458^DAFSYG^463^ in *Pc*POx H158Y, the hydrogen binding between Thr160 and Asp458 suggested it was a kind of sugar binding mode, since the rotation of the threonine residue and forming a hydrogen bond to aspartic acid was a trigger of substrate binding in *To*POx [41]. The orientation of the glycerol molecule in the catalytic centre, which showed a good fit with the orientation of the sugar substrate analog, also suggested this view. The orientation of the glycerol lacked that of the C6 hydroxymethyl group of 3-fluoro-3-deoxy-D-glucose. Considering Tyr462 in the loop ^458^DAFSYG^463^ is responsible for recognition of the C6 hydroxymethyl group [41], the lack of C6 orientation of the glycerol may cause missing the loop. Only Asp458, which forms a hydrogen bond to Thr160, is fixed and others in the loop ^458^DAFSYG^463^ are flexible due to missing recognition sites. In case of *Op*POx, the crystal structure (PDB ID: 9FL2) without sugar substrates is in closed conformation despite the presence of glycerol as cryoprotectant [48]. The absence of the glycerol molecule in the *Op*POx active site can be explained by the sensitive substrate recognition of *Op*POx. The *K*_m_ value for glucose is 0.048 mM, much lower than that of *Pc*POx (0.88 mM).

To form a covalent flavin attachment, at least a suitable conformation of the flavin and activation of the corresponding residue are necessary. In particular, hydrogen bonding by Thr169 and a positive charge of the catalytic base His548 are critical for covalent flavin formation (quinone methide mechanism) in *To*POx [35]. Including these two residues, there is no substantial conformational difference in the environment around FAD between *Pc*POx WT and H158Y, except for the residue His158 or Tyr158. The aromatic side chain of Tyr158 in the *Pc*POx H158Y mutant altered its conformation compared to the imidazole ring of His158 in *Pc*POx WT, making the covalent interaction to FAD unavailable. This conformational change can also be observed in the corresponding histidine residues of POx and C-glycoside oxidase (CGOx), which share a common ancestor. [25,48,52,54–57]. Although POxs form covalent flavin interactions with histidine residues, CGOxs do not form a covalent flavin bond, even though the histidine residues are conserved. The crystal structures of POx and CGOx are compared in Fig. S5. The conformational change of the histidine prevents forming a covalent interaction to FAD in CGOx, as well as in the *Pc*POx H158Y mutant. The noncovalent mutant T169A (PDB ID: 3LSH) has a relatively similar orientation of His167 (site linked to FAD) to that of WT, but a hydrogen bond between His167 and Thr158 still fixes His167 in a wrong conformation to form a covalent flavin bond. *To*POx WT shows a Asp81-Lys90-Gln110-His167 network, but the noncovalent mutant T169A loses this network [35]. These informations suggest the histidine/tyrosine conformation observed in *Pc*POx H158Y or CGOx is an initial orientation of the aromatic side chain before covalent flavin formation; His167 in *To*POx T169A is an intermediate conformation during formation of covalent flavin, which is still not able to form a covalent FAD, bond, and the histidine linked to FAD observed in various POxs is the final evolutionary conformation (Fig. 7).

**Fig. 7.**
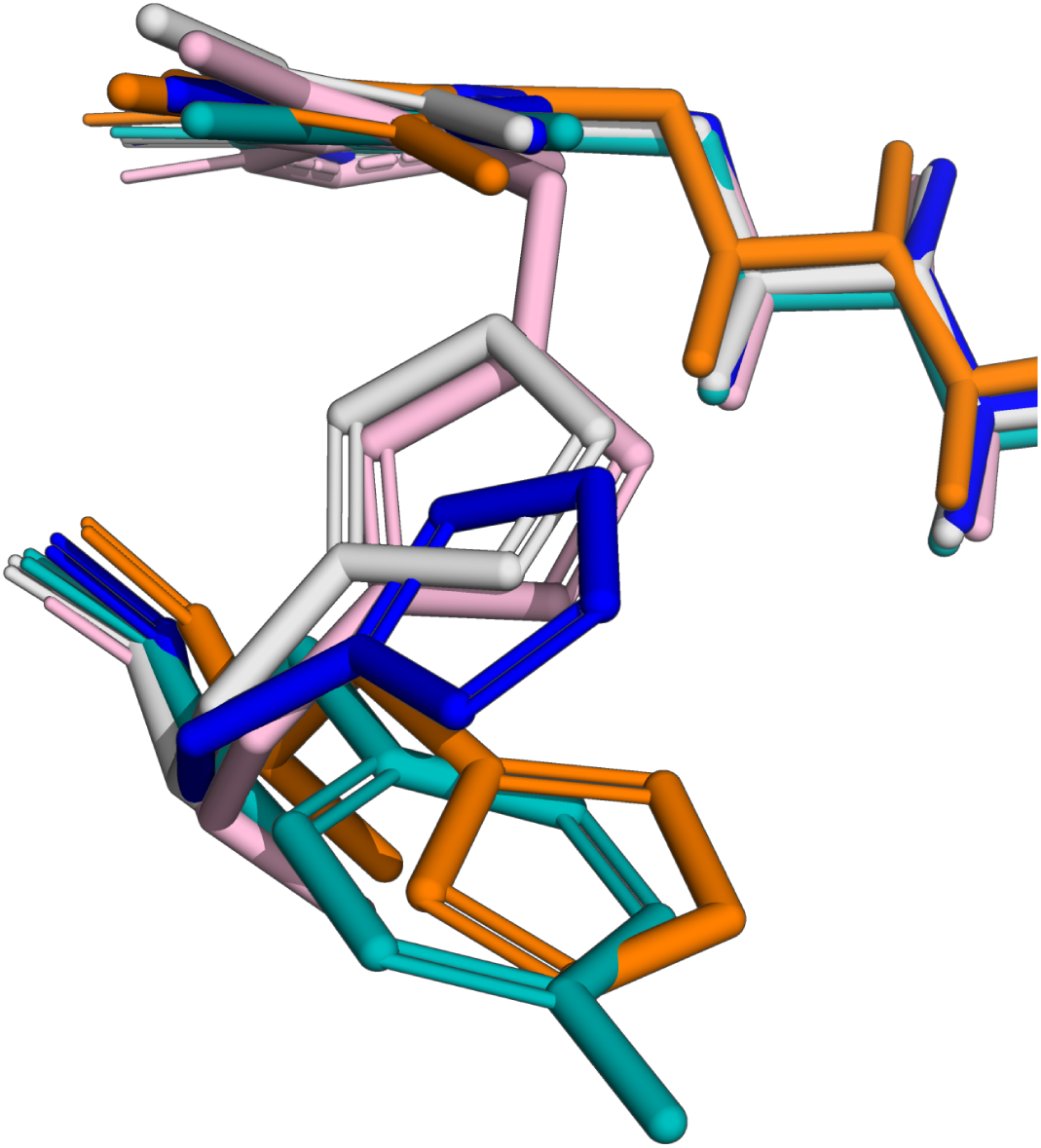
Conformation of Tyr158 in *Pc*POx H158Y and a corresponding histidine residue in *Pc*POx WT (PDB ID: 4MIG), *To*POx WT (PDB ID: 1TT0), *To*POx T169A (PDB ID: 3LSH), and *C*-glycoside oxidase from *Pseudarthrobacter siccitolerans* (*Ps*CGOx), WT (PDB ID: 7QF8). *Pc*POx WT (magenta) and *To*POx WT (gray) are connected with FAD. *To*POx T169A (dark blue), *Ps*CGOx (orange), and *Pc*POx H158Y (cyan) are not connected with FAD.

We introduced the mutation K79A in the *Pc*POx H158Y background to release the hydrogen bond between Lys79 and Tyr158, expecting a conformational change of Tyr158. The double mutant H158Y/K79A did not form a covalent FAD bond, but whether Tyr158 altered its conformation in the double mutant is still not known. In addition to the proper conformation, deprotonation of the phenolic hydroxyl group of Tyr158 is essential for its nucleophilic attack to C8α of FAD in the proposed autocatalytic mechanism of covalent flavin attachment [58]. The high p*K*_a_ value of Tyr158 in *Pc*POx H158Y (8.96) made covalent flavin formation difficult in neutral conditions. In case of *p*-cresol methylhydoxylase (PDB ID: 1DII), which has an 8α-*O*-tyrosyl covalent flavin linkage [4], a carboxylate oxygen of Asp440 is located close to Oη of Tyr158 (5.4 Å) (Fig. S6). This Asp440-Tyr384 interaction can assist deprotonation of the phenolic hydroxyl group of Tyr384 before covalent flavin formation. To form 8α-*O*-tyrosyl covalent flavin bond in *Pc*POx, not only the tyrosine conformation but also a residue decreasing the p*K*_a_ value of Tyr158 may be necessary. Overall, the reason for the disconnection of Tyr158 to FAD is mainly due to the conformation of the side chain of Tyr158, and the K79A mutation fails to restore a favorable conformation or rescue the covalent flavin formation.

The H158Y mutation decreased the oxidase activity for all tested sugar substrates. The *Pc*POx H158Y mutant showed comparable *K*_m_ values for xylose and 2-deoxy-D-glucose, suggesting it is not compromised substrate recognition that causes the decrease of activity. The H158Y mutation also significantly decreased the dehydrogenase activity for both DCIP and BQ. As well as many sugar oxidoreductases, POx undergoes mainly two half reactions: a reductive half reaction (sugar oxidation/flavin reduction) and an oxidative half reaction (oxygen reduction/flavin oxidation) [59]. Our results suggested that the reductive half reaction decreased due to the H158Y mutation since the decrease of enzymatic activity was independent of the electron acceptors. The previous pre-steady kinetic research on *To*POx H167A variant revealed that the loss of the covalent flavinylation critically affected the reductive half reaction, but not the oxidative half reaction [60]. A reason for the impaired reductive-half reaction may be a lowered redox potential; the redox potential value of *To*POx WT is −115 mV and that of *To*POx H167A is −147 mV [24]. The decrease in enzymatic activity in *To*POx is not simply attributable to substitutuion of His167, because *To*POx T169A variant, which retains His167 but contains non-covalently bound flavin, also shows a marked reduction in catalytic turnover (the *k*_cat_ values for glucose in WT and H169A mutant were 152 s^−1^ and 0.69 s^−1^ respectively) [61]. These observation suggests a functional relationship among the covalent flavin linkage, the reductive half reaction, and redox potential in *To*POx. A similar situation has been seen in pyranose dehydrogenase from *Agaricus meleagris* (*Am*PDH). The noncovalent variant *Am*PDH H103Y shows a significantly slower reductive-half reaction but maintains the rate of the oxidative half-reaction compared to *Am*PDH WT [62]. A decrease in the redox potential is also observed in *Am*PDH H103A mutant [63]. Therefore, the lower turnover numbers of *Pc*POx H158Y are assumed to be due to a slower reductive half reaction, caused by lower redox potential from the loss of covalent flavin linkage. To verify this hypothesis, the measurement of the reductive half reaction, the oxidative half reaction, and the redox potential of *Pc*POx and H158Y mutant are required.

On the sugar substrate recognition of *Pc*POx H158Y mutant, changes in *K*_m_ values tend to be more pronounced for substrates with lower *K*_m_ values, such as glucose and galactose. Structural observations may explain the reason: there was no significant difference except for the covalently linked flavin cofactor in the structures of the catalytic centres of *Pc*POx WT and the H158Y mutant, implying the recognition of only sensitive substrates can be affected by H158Y mutation. For the recognition of oxygen molecules in the oxidative half reaction, the catalytic pair His and Asn/His, which is widely conserved in CAZy AA3 sugar oxidoreductases is considered important since the catalytic pair recognizes oxygen molecule close to the *re*-face of flavin in glucose oxidase [64]. This insight agrees with the conversion from oxidase to dehydrogenase in *To*POx caused by the N593C mutation [65,66]. In the case of *Pc*POx, the catalytic pair His553 and Asn596 does not change its conformation by the mutations H158A or H158Y, partly explaining the consistent oxidative half reaction rates among these variants.

In conclusion, the *Pc*POx H158Y structure demonstrates that tyrosine substitution at the flavinylation site does not support covalent flavinylation primarily because Tyr158 adopts an unproductive rotamer conformation that places its phenolic oxygen too far from FAD C8α. Disruption of the Tyr158-Lys79 hydrogen bond, which stabilizes this unproductive conformation, by the mutation K79A did not restore covalent flavinylation, indicating that additional constraints prevent formation of an 8α-*O*-tyrosyl flavin. Loss of covalent flavinylation markedly decreased both oxidase and dehydrogenase turnover rates, likely due to impairment of the reductive half-reaction. These findings provide a structural explanation for the failure of tyrosine substitution to enable covalent flavinylation in *Pc*POx and highlight key requirements for engineering alternative covalent flavin linkages in covalent flavoproteins.

## Abbreviations

Pox: pyranose oxidase
*Pc*POx: pyranose oxidase from *Phanerochaete chrysosporium*
CGOx: C-glycoside oxidase

## Acknowledgements

This study received financial support from Grants-in-Aid for Scientific Research (A) from the Japan Society for the Promotion of Science (JSPS) (No. 23H00341 to KI) and a Grant-in-Aid for JSPS Fellows (No. 24KJ0866 to YY).

## Data accessibility

The structural data of *Pc*POx H158Y mutant are openly available in the Protein Data Bank under accession code: 9X0E.

## Author contributions

KI supervised the study; YY and CKP designed experiments; YY performed experiments; YY analyzed data and wrote the first draft of the manuscript, TU and KT revised the manuscript. All authors made manuscript revisions. All authors read and approved the final version of the manuscript.

