## Supplementary material for "Structural basis for loss of covalent flavinylation in the H158Y mutant of pyranose oxidase": 260217YashimaSuppl.pdf

**Table S1.** Primers for the preparation of the double mutant K79A and H158Y.

| Mutation | Primer | Sequence |
|---|---|---|
| K79A | Forward | 5´-TATCATGCGAAGAACGAGATCGAGTACCAGAAGGACATCG-3´ |
| K79A | Reverse | 5´-GTTCTTCGCATGATACCCAGGGATGGGAACCTG-3´ |
| H158Y | Forward | 5´-AGCACTTACTGGACGTGCGCAACCCCCGAG-3´ |
| H158Y | Reverse | 5´-CGTCCAGTAAGTGCTCATGCCCCCGACGCC-3´ |

**Table S2.** Comparison of *Pc*POx codon defferences in this study (PDB ID: 9X0E) and the previous structure (PDB ID: 4MIF). Differences in codon are highlighted in red. For amino acid residues, silent mutations are shown in black, and amino acid substitutions are highlighted in red.

| | PDB ID: 9X0E | PDB ID: 4MIF* |
|---|---|---|
| Residue number | Codon | Codon |
| 9 | AGT (Ser) | CGT (Arg) |
| 132 | GGA (Gly) | GGC (Gly) |
| 167 | TTC (Phe) | TTT (Phe) |
| 356 | CCT (Pro) | CCC (Pro) |
| 359 | GAA (Glu) | GAG (Glu) |
| 362 | TCT (Ser) | TCG (Ser) |
| 367 | CGT (Arg) | CGC (Arg) |
| 441 | GTA (Val) | GTC (Val) |
| 483 | ACA (Thr) | ACC (Thr) |
| 507 | ACG (Thr) | ACA (Thr) |
| 509 | AAC (Asn) | AAG (Lys) |
| 520 | GCT (Ala) | GCC (Ala) |
| 525 | GAC (Asp) | GAT (Asp) |

*The nucleotide sequence was obtained from NCBI accession number AY522922. 1. The corresponding amino acid sequence for AY522922. 1 is identical to that of the crystal structure of *Pc*POx WT (PDB ID 4MIF).

**Table S3.** The predicted p$K_a$ values of the tyrosine158 and the lysine79 in the crystal structure of *Pc*POx H158Y mutant.

| | p$K_a$ | Locate | Desolvation effects | | | | Sidechain | | Backbone | | Coulombic Interaction | |
|---|---|---|---|---|---|---|---|---|---|---|---|---|
| | | | Massive | | Local | | Hydrogen bond | | Hydrogen bond | | | |
| Tyr158 | 8.96 | BURIED | 0.46 | 446 | 0.14 | 2 | -0.32<br>-0.79 | Thr157<br>Lys79 | 0.00000<br>0.00000 | 0<br>0 | 1.87<br>-2.40 | Asp48<br>Lys79 |
| Lys79 | 14.61 | BURIED | -0.27 | 427 | -0.42 | 6 | 0.00000<br>0.00000 | 0<br>0 | 0.00000<br>0.00000 | 0<br>0 | 2.40<br>2.40 | Asp48<br>Tyr158 |

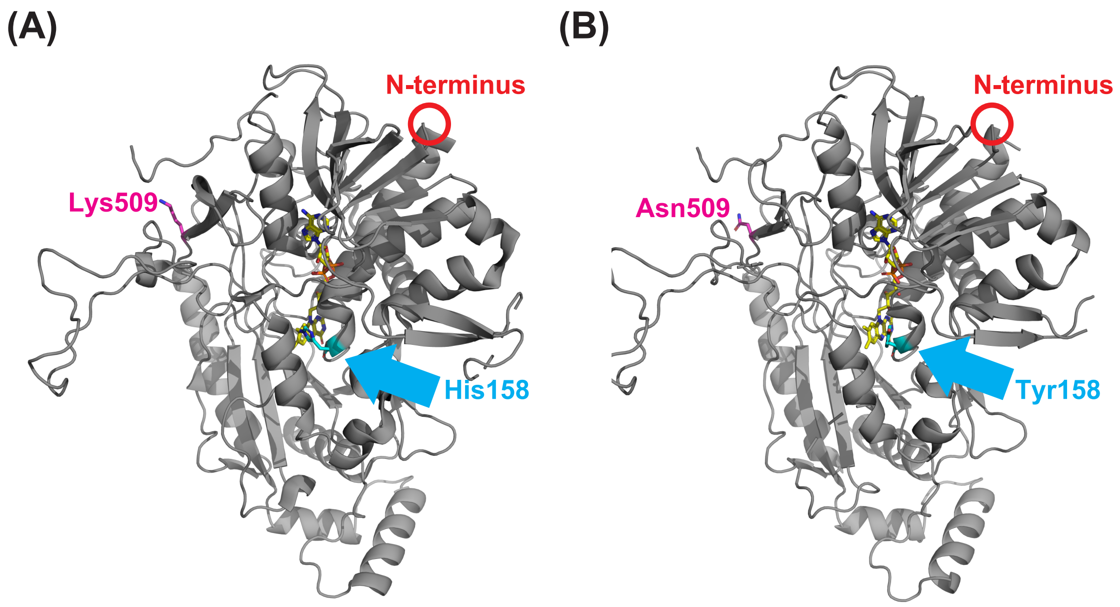


**Fig. S1.** The structure of *Pc*POx in this work compared with the previous work. Our *Pc*POx we cloned and used in this study contained two amino acid substitutions R9S and K509N compared to the *Pc*POx WT reported previously (PDB ID: 4MIG). (A) The overall monomeric structure of *Pc*POx WY (PDB ID: 4MIG). Red circles indicate the edges of the N-terminal Pro13 since the structure does not show N-terminus including Arg9. The residues Lys509, His158, and FAD cofactor are shown in magenta, blue, and yellow respectively. (B) The overall monomeric structure of *Pc*POx H158Y mutant (PDB ID: 9X0E). Red circle indicates the edges of the N-terminal Pro13 due to the same reason as (A). The residues Lys509, Tyr158, and FAD cofactor are shown in magenta, blue, and yellow respectively.

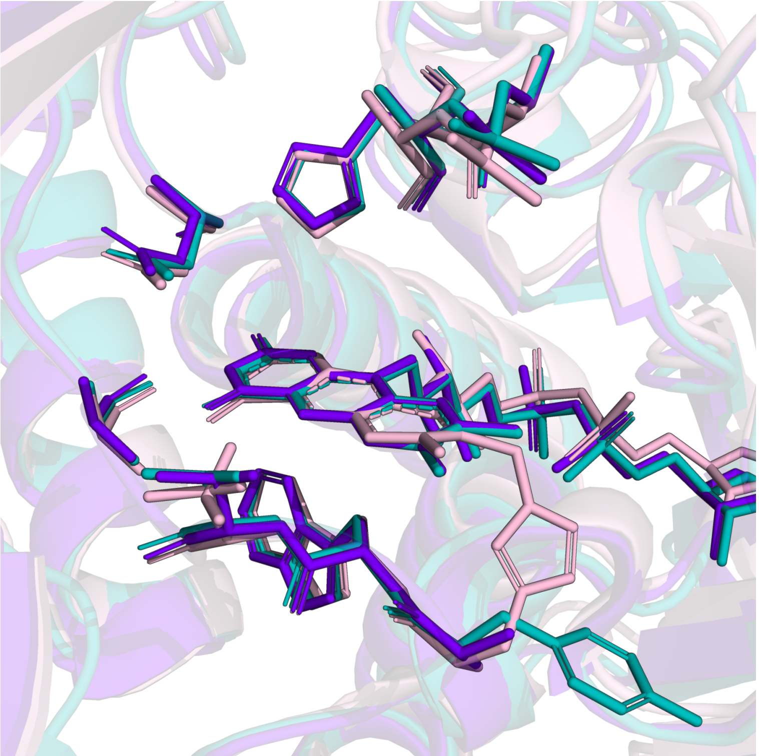

**Fig. S2.** Comparison of the amino acids around the isoalloxazine ring of FAD cofactor in *Pc*POx WT (PDB ID: 4MIG), *Pc*POx H158A (PDB ID: 4MIH), and H158Y(PDB ID: 9X0E). The protein backbone is shown in magenta (WT), purple (H158A) and cyan (H158Y).

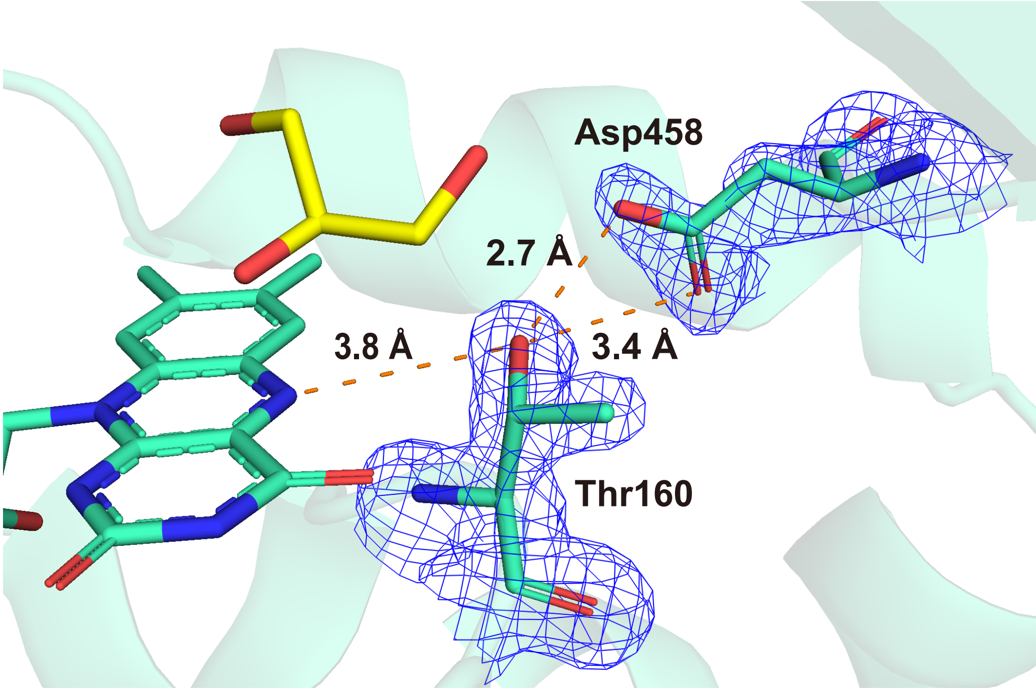


**Fig. S3.** The conformation of  Thr160 and hydrogen bond interaction to Asp458.

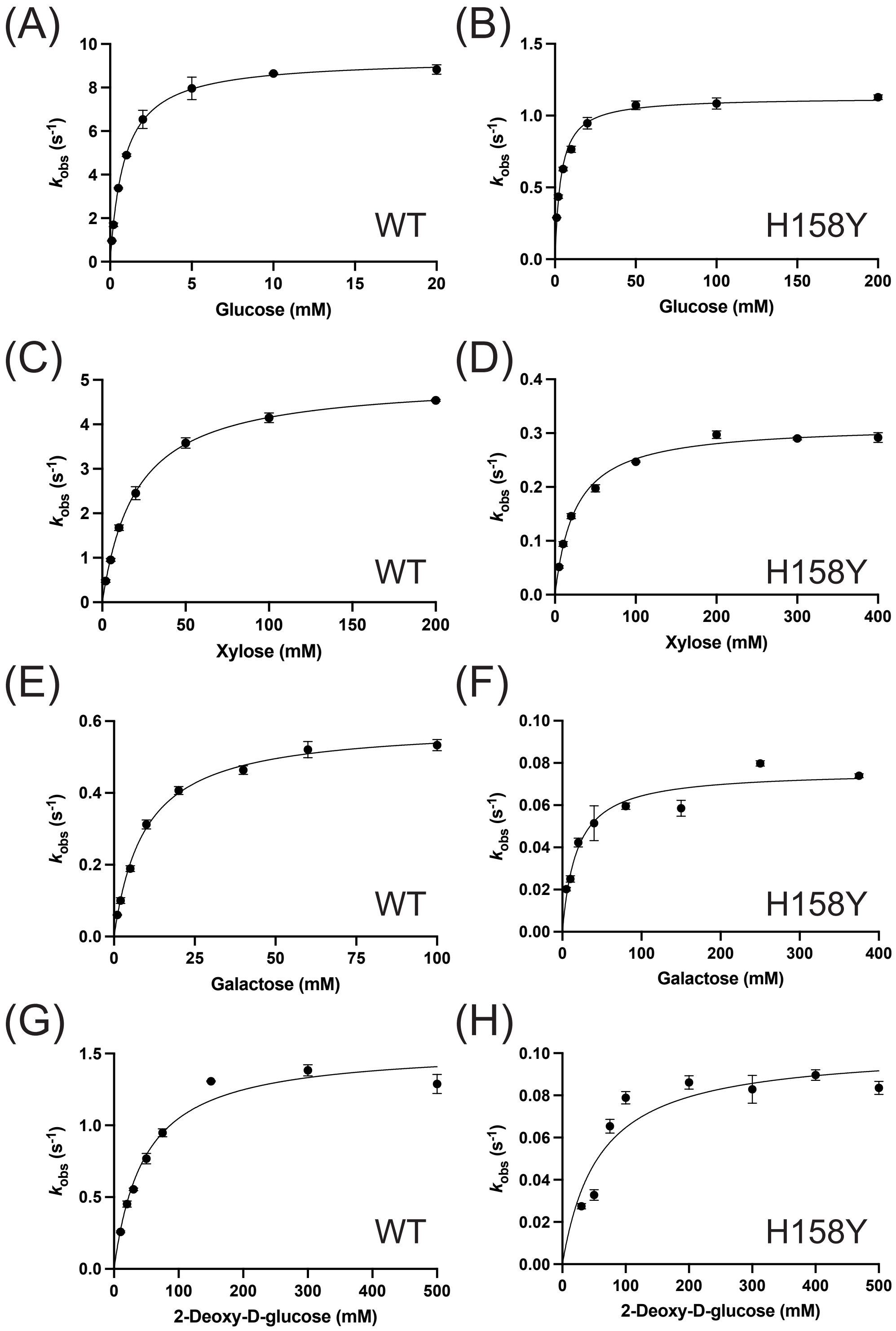


**Fig. S4.** Michaelis-Menten curve fitting of *Pc*POx WT and the H158Y variant with the tested sugar substrates. Error bar indicates the SD calculated from data collected in triplicate ($n = 3$).

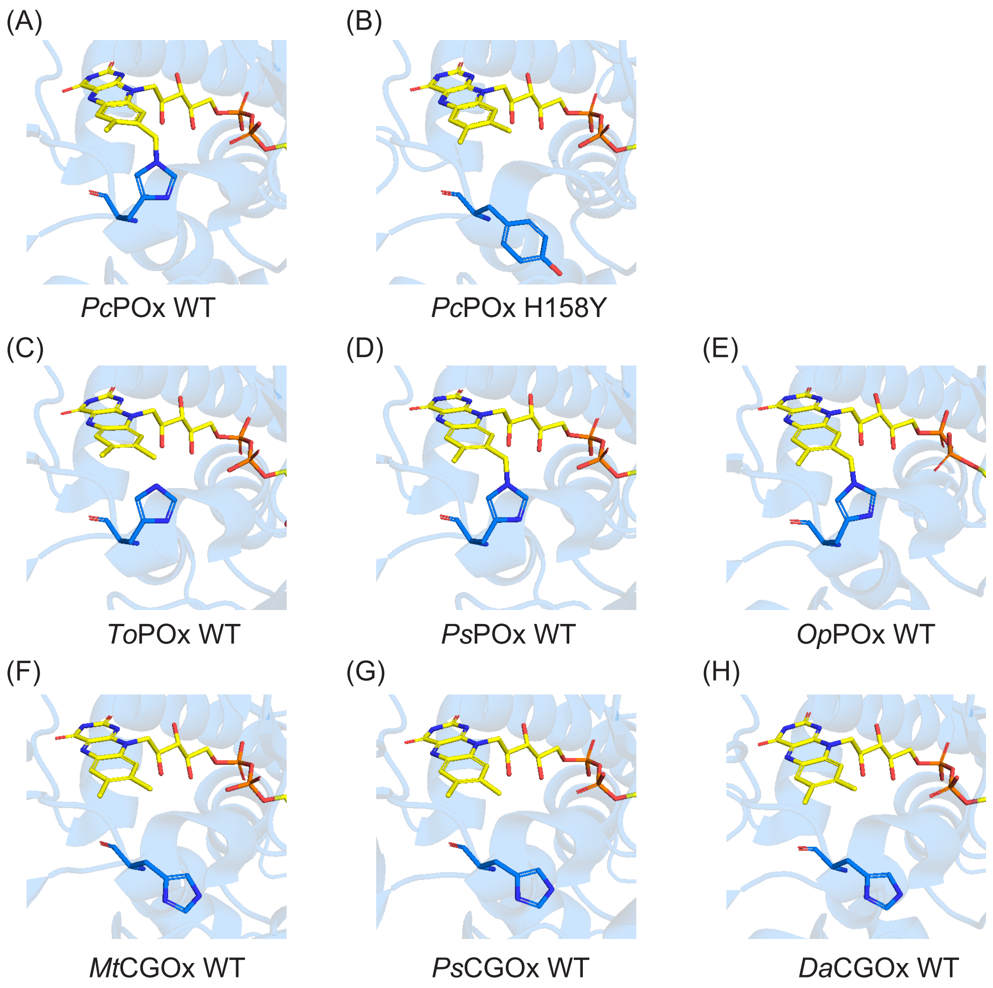


**Fig. S5.** Interaction between FAD and histidine residues in pyranose oxidases and C-glycoside oxidases. (A) Pyranose oxidase from *Phanerochaete chrysosporium*, WT (PDB ID: 4MIG). (B) Pyranose oxidase from *Phanerochaete chrysosporium*, H158Y (PDB ID: 9X0E). (C) Pyranose oxidase from *Trametes ochracea*, WT (PDB ID: 1TT0). (D) Pyranose oxidase from *Peniophora* sp., WT (PDB ID: 1TZL). (E) Pyranose oxidase from *Oscillatoria princeps* RMCB-10, WT (PDB ID: 9FL2). (F) *C*-glycoside oxidase from *Microbacterium trichothecenolyticum*, WT (PDB ID: 7DVE). (G) *C*-glycoside oxidase from *Pseudarthrobacter siccitolerans*, WT (PDB ID: 7QF8). (H) *C*-glycoside oxidase from *Deinococcus aerius*, WT (PDB ID: 8QVE). Structures (A), (C), (D), and (E) form a covalent bond between FAD and the corresponding histidine residue; however, the covalent FAD linkage is not graphically shown in the *To*POx structure (C). In contrast, the other structures do not form a covalent flavin.

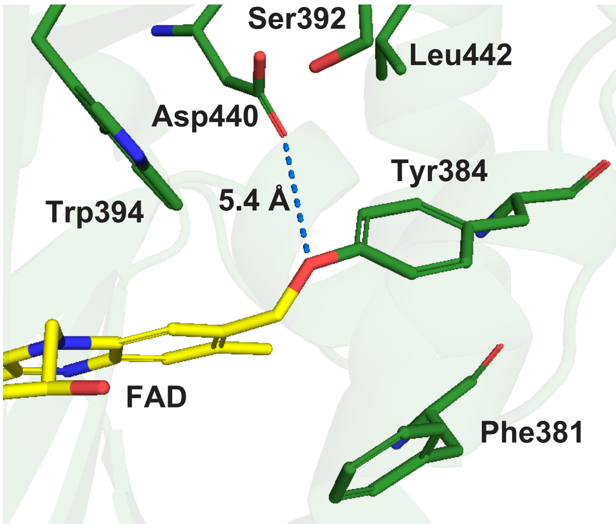


**Fig. S6.** The covalent interaction between FAD cofactor and Tyr384 in *p*-cresol methylhydoxylase (PDB ID: 1DII) Supplementary Materials for
